# Flotillin acts as a regulatory cellular barrier against invasion by *Escherichia coli* K1

**DOI:** 10.64898/2026.10.27.762333

**Authors:** Lip Nam Loh, Andrea Zelmer, Theresa H. Ward

**Affiliations:** Department of Infection Biology, Faculty of Infectious and Tropical Diseases London School of Hygiene & Tropical Medicine London U.K

## Abstract

Pathogenic microorganisms that cause neonatal meningitis must cross the blood-brain barrier to reach the central nervous system. *Escherichia coli* K1 remains a leading pathogen in cases of neonatal bacterial meningitis and invades the endothelium in the brain microvasculature. Signalling through a cascade of sphingosine kinase activity and the epidermal growth factor receptor (EGFR) has been implicated in this process, but how it is coupled to membrane organisation is unclear. Here we show that invasion of human brain microvascular endothelial cells by *E. coli* K1 can also be regulated by flotillins through the sphingosine/EGFR axis. Loss of flotillin 1 markedly increased bacterial invasion, whereas flotillin overexpression suppressed it, and the hyperinvasive phenotype of flotillin-deficient cells remained dependent on cholesterol. Depletion of sphingosine kinases by siRNA also increased invasion, in contrast to pharmacological inhibition, suggesting that the kinase proteins, and not only their catalytic activity, may help restrain bacterial entry. In a neonatal mouse infection model, flotillin 1 knockout animals had lower bacteraemia but equivalent brain bacterial burdens, indicating increased penetration of the blood-brain barrier. We propose that flotillin-dependent membrane organisation restrains an EGFR- and sphingolipid-dependent route of *E. coli* K1 entry, defining a host barrier to meningitic infection.

## Introduction

Bacterial meningits remains a global threat to health and *Escherichia coli* K1 is a leading cause of neonatal bacterial meningitis, a disease that continues to carry substantial mortality and a high risk of long-term neurological sequelae. Once a threshold blood bacteraemia is reached^1^, circulating bacteria cross the blood-brain barrier (BBB), and invasion of brain microvascular endothelial cells is a key step in this process^2^. The host pathways that *E. coli* K1 exploits are still incompletely defined, and rising antimicrobial resistance has increased interest in host-directed approaches that could complement antibiotics.

Recent work has identified a signalling route used by meningitic *E. coli*. Wang et al.^3^ showed that bacterial OmpA, FimH and NlpI activate sphingosine kinase 2 (SphK2) in human brain microvascular endothelial cells (HBMEC), and that the resulting sphingosine-1-phosphate (S1P) signals through S1P receptor 2 to activate the epidermal growth factor receptor (EGFR) and its downstream kinase c-Src. Bacterial entry also depends on membrane cholesterol, and *E. coli* K1 invades through an actin-dependent, macropinocytic route that requires intact lipid rafts^4^. How this signalling is coupled to membrane organisation at the cell surface of the endothelial cells is not known.

Flotillin 1 and flotillin 2 are good candidates for this link. They co-assemble into cholesterol-dependent microdomains at the plasma membrane and on late endosomes, and define a clathrin-independent endocytic pathway^5^. Furthermore, flotillin has been implicated in down regulating EGFR signaling and thereby controlling EGFR-dependent micropinocytosis^6,7^. Flotillins also bind sphingosine, recruit it to membranes and thereby maintain cellular S1P levels^8^, which raises the possibility that they might be involved in the same sphingolipid pathway implicated in *E. coli* K1 invasion.

Here, we examined the EGFR and sphingosine kinase pathway in HBMEC and investigated how flotillin might modulate *E. coli* K1 invasion. We show that *E. coli* K1 invasion depends on EGFR, sphingosine kinase activity and cholesterol-rich membrane domains, that flotillin 1 associates with bacteria during infection, and that loss of flotillin enhances invasion *in vitro* and increases bacterial penetration into the brain in a neonatal mouse model. Together, these data identify flotillin as a cholesterol-dependent restraint on *E. coli* K1 entry across the BBB.

## Materials and Methods

### Cell culture and bacteria

Details of cells and strains used were provided^4^. Additionally, *E. coli* K1 (A192PP), a gift from Peter Taylor, UCL, was utilized for *in vivo* infections^1^.

### Antibodies

Antibodies used were: monoclonal mouse anti-b-actin (clone AC-15, Sigma), Alexa Fluor 488-labelled monoclonal EGFR antibody (Thermo Fisher Scientific), mouse anti flotillin 1 (BD Biosciences), rabbit polyclonal anti-sphingosine kinase 1 (St John’s Laboratory), rabbit polyclonal anti-sphingosine kinase 2 (St John’s Laboratory), polyclonal rabbit antibody to *E. coli* O18 (Mast Group), FluoProbes-secondary antibodies (Interchim, Cheshire Science Ltd.), horseradish peroxidase (HRP)-conjugated goat anti-mouse and HRP-conjugated goat anti-rabbit (Bio-Rad).

### Transient transfection of HBMEC

Plasmids used in this study were as follows: EGFR-GFP WT (Addgene 32751) and perbB1Y1045F-EGFP (Addgene 40267), p58-GFP^9^, the SphK1 plasmids were a kind gift from Stuart Pitson (University of South Australia, Adelaide) and included pCDNA3.1, SphK1-GFP WT^10^, SphK1-GFP G113A^10^, SphK1-GFP G82D^11^; AcGFP-C2-hSphK2 (Addgene 84370), pSphK2 WT (Addgene 20571), LDLR A18-GFP (G.E. Kreitzer, University of Cornell)^12^, caveolin-1-GFP^13^, flotillin 1-GFP^5^, flotillin 2-GFP^14^.

For plasmid transfection, HBMEC were seeded onto glass coverslips, grown to approx. 80% confluency and then transfected with jetPRIME reagent (Polyplus-transfection SA) according to the manufacturer’s instructions. Infection with *E. coli* K1 was undertake approx 20 hours post-transfection.

### siRNA Transfection

Non-targeting siRNA or sphk1, sphk2 or flotillin 1 siRNA were all SMARTpool siGENOME siRNA from Dharmacon, GE Healthcare). These were delivered to cells at a final concentration of 50 µM, using Lipofectamine RNAiMAX (Thermo Fisher), following the manufacturer’s protocol. Assays were carried out 48 hours after transfection.

### Invasion assay

Following transfection, the invasion assay was conducted as per Loh et al^4^. In brief, HBMEC were grown to confluency, with triplicate wells for each condition.

Experiments were conducted in low serum media, and pretreated as described for each experiment. Infection was with moi of approximately 100 with mid-log phase *E. coli* K1 or K-12. Cells were incubated at 37°C as indicated, after which cells were overlaid with growth media containing 100 µg/ml gentamicin for 1 hour at 37°C to kill extracellular bacteria. Cells were trypsinised and an aliquot retained for cell counting, before being lysed with 0.3% SDS in PBS. Bacteria cfu assessed by titration and cells were counted on a haemocytometer. Results were expressed as the number of intracellular bacteria per cell (measured by cfu/total cell number per well) relative to control. At least 3 independent experiments were undertaken.

### Pharmacological inhibitors

Cells were pretreated for 30 min with reagents or control vehicle in experimental media, prior to infection as above with *E. coli* K1 in continuing presence of the reagent. Gefitinib (DMSO, Sigma) was used at 10 µM, erlotinib (DMSO, Sigma) was used at 10 µM, PF543 (water, Pfizer) was used at 10 µM, ABC294640 (water, Active Biochemicals) was used at 50 µM, SKI II (DMSO, Sigma) was used at 20 µM. Control experiments with methyl-β-cyclodextrin (MβCD, Sigma) were conducted in serum-free media with MβCD at 5 µM. Reagents were tested for potential toxicity on HBMEC by trypan blue exclusion, and their effects on bacterial viability were tested by cfu analysis.

### Invasion assay

Invasion assay was conducted as per Loh et al^4^. In brief, HBMEC were grown to confluency, with triplicate wells for each condition. Experiments were conducted in low serum media, and pretreated as described for each experiment. Infection was with moi of approximately 100 with mid-log phase *E. coli* K1 or K-12. Cells were incubated as indicated, after which cells were overlaid with growth media containing 100 µg/ml gentamicin for 1 hour at 37°C to kill extracellular bacteria. Cells were trypsinised and an aliquot retained for cell counting, before being lysed with 0.3% SDS in PBS. Bacteria cfu assessed by titration and cells were counted on a haemocytometer. Results were expressed as the number of intracellular bacteria per cell (measured by cfu/total cell number per well) relative to control. At least 3 independent experiments were undertaken.

### Immunofluorescence

HBMEC cells grown on coverslips were transfected and infected as above. Following incubation with bacteria at 37°C, cells were fixed in 3% formaldehyde/PBS for 15 min at room temperature, washed with PBS and then incubated for 10 min in 50 mM NH_4_Cl in PBS. Cells were washed in 5% FCS/PBS and then labelled sequentially with primary and secondary antibodies diluted in wash buffer with 0.2% saponin, each with 1 hour room temperature incubations.

For differential staining of intra- and extra cellular bacteria, cells were initially stained with anti-*E. coli* O18 without permeabilization, and then FluoProbes-642 donkey anti rabbit IgG, before a second sequential staining step in the presence of saponin with anti-*E. coli* O18 in combination with other antibodies as appropriate. Following this second stage with primary antibody incubation, a second fluorophore was used to detect the bacteria, usually FluoProbes-546. Cells were mounted with Fluoromount-G containing DAPI (eBioscience).

Samples were observed with an inverted Zeiss LSM 880 confocal laser-scanning microscope with appropriate lasers. Images were processed with ImageJ and Adobe Photoshop.

### Flow cytometry

HBMEC cells were infected as above, then trypsinised. Once the cells were in suspension they were stained with AF488-labelled anti-EGFR at 1:100 for 15 min on ice. Cells were washed and suspended in FACS buffer (1% FBS in PBS).

Acquisiion of flow cytometry data was performed on a BD LSRII (BD Biosciences) using appropriate laswer lines. Data was analysed using FlowJo software (version 10, Tree Star, Ashland). Gating around cells was quantified for median fluorescence in the AF488 channel. Average background was subtracted from the MFI and relative fluorescence was quantified against the positive uninfected control.

### Animals

All animal studies were conducted under UK Home Office license 70/8603 and were approved by the LSHTM Animal Welfare and Ethical Review Board.

C57 BL/6 mice and *flotillin-1^-/-^* adult mice^8^ were time-mated using Whitten effect to synchronise litters. Male and female pups from 3-4 litters were used for experimental infection at day 5-7.

#### Murine infection model

10 µl *E. coli* K1 A192PP culture (5 X 10^6^ bacteria/ml resuspended in saline solution) were injected intraperitoneally into neonatal pups at age d5-7. After bacterial administration, mouse pups were monitored at regular intervals and sacrificed when symptoms progressed to defined stages according to a symptom score chart or at 24 hours whichever was the sooner. Blood and brain tissue were collected by humane procedures. Brain tissue was placed in PBS on ice, blood was deposited in tubes containing 5 µl heparin.

#### Bacterial burdens

Brain was homogenized by pushing through a tissue sieve. Brain homogenate and blood were serially diluted in PBS, and bacteria were enumerated by standard plate counting on LB agar. Agar plates were incubated at 37°C overnight.

### Statistical analysis

All experiments were repeated 3 times and performed in triplicate. Data are expressed as denoted in figure legends. Bacterial counts from tissue were log-transformed before analysis, and analysed by unpaired *t*-tests and one-way analysis of variance (with *post hoc* Bonferroni tests). All statistical analyses were performed with GraphPad Prism software. *p* < 0.05 was considered statistically significant.

## Results

### *E. coli* K1 utilises EGFR and the sphingosine kinase pathway to invade HBMECs

Building on the findings of Wang et al.^3^, who demonstrated that S1P-mediated activation of EGFR is exploited by meningitic *E. coli* for BBB penetration, we investigated whether this pathway operates in our HBMEC model using complementary genetic and pharmacological approaches.

We firstly looked at the role of EGFR in *E. coli* K1 invasion though overexpression of GFP-tagged EGFR chimaeras. Overexpression of wild-type GFP-tagged EGFR significantly increased *E. coli* K1-infected cells to approximately 200% relative to the transfection control, whilst expression of the dominant active Y1045F mutant^15^ produced a further significant increase to approximately 320% (Fig. 1A). Next, cells were pretreated with the EGFR inhibitors gefitinib or erlotinib prior to infection.

**Figure 1.**
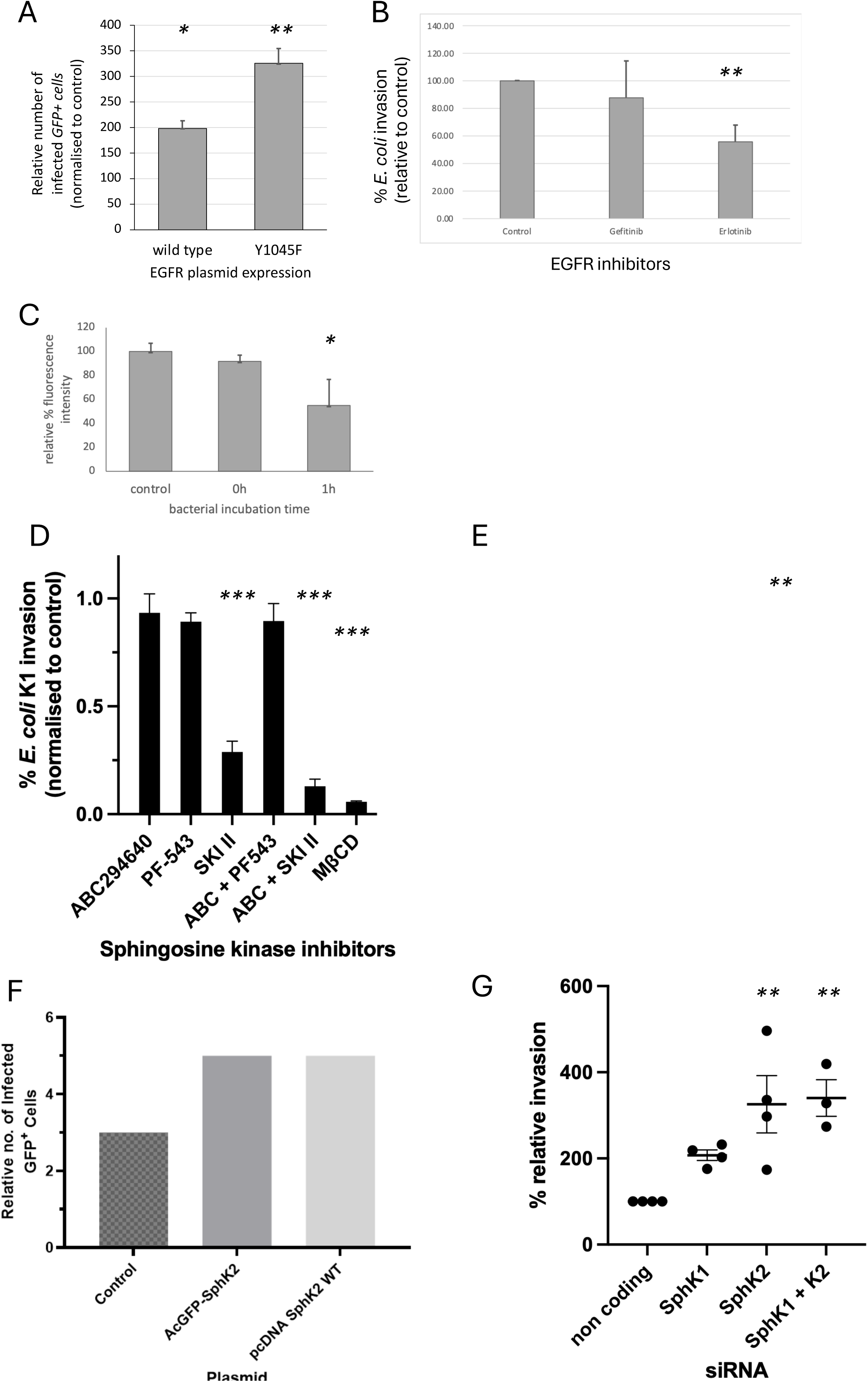
*E. coli* K1 uses EGFR and the sphingosine kinase pathway for invasion into HBMEC. A. Uptake of E. coli K1 following transfection of HBMEC with GFP-EGFR plasmids. GFP-expressing cells were scored for the presence of intracellular *E. coli* K1 following infection at moi 100 for 2 hours. The data is normalised to the transfection control cells expressing a membrane-bound GFP chimaera (p58-GFP a membrane protein that localises to the early secretory pathway and is not involved in plasma membrane dynamics). The left column shows GFP-tagged WT EGFR and the right column shows GFP-tagged Y1045F EGFR, a dominant positive mutant. Both show a significantly positive increase over the control cells, with the positive mutant facilitating a further increase over wild type. B. Cells were pretreated with EGFR-inhibitory agents for 30 min, then infected with *E. coli* K1 for 1 hour. Internalised bacteria were assayed by colony counting and normalised to mock treatment. C. HBMEC were infected with *E. coli* K1 for 1h, 0h (bacteria added then washed away) or uninfected. Cells were trypsinised and stained with Alexa-Fluor 488 labelled anti-EGFR on ice. Surface labelling of EGFR was analysed by flow cytometry. D. HBMEC were pretreated with Sph kinase inhibitors for 30 minutes, then infected with *E. coli* K1 for 1 hour. Internalised bacteria were assayed by colony counting and normalised to the appropriate mock treatment, either RPMI alone or with DMSO. E. HBMEC were transiently transfected with SphK1-GFP chimaeras: WT or gain of function mutant G113A or kinase dead G82D mutant as indicated. pCDNA3.1 was a transfection control. Cells were infected with *E. coli* K1 for 2h, then internalised bacteria were assayed by colony counting and normalised to the control. F. HBMEC were transiently transfected with two wild type SphK2-GFP chimaeras to investigate overexpression effects. GFP-expressing cells were scored for the presence of intracellular *E. coli* K1 following infection for 2 hours. G. Uptake of *E. coli* K1 for 2 hours following siRNA depletion of Sph kinases. Relative invasion was determined against noncoding siRNA control. Data shown are mean ± SD from >3 independent experiments with triplicate samples. * *p*<0.05, ** *p*<0.01, *** *p*<0.005

Gefitinib reduced invasion to approximately 88% of control, though this did not reach statistical significance, whilst erlotinib produced a more pronounced and significant reduction to approximately 53% of control (Fig. 1B). By utilising a fluorophore-tagged EGFR antibody, changes to surface expression of EGFR could be measured.

When *E. coli* K1 were introduced to the cells and incubated for one hour, surface expression of EGFR was significantly reduced compared to uninfected controls, suggesting receptor internalisation upon bacterial engagement, whilst addition of bacteria followed by immediate washing (0h) had no significant effect (Fig. 1C). These results were consistent with EGFR involvement in *E. coli* K1 uptake into HBMEC.

The role of the sphingosine kinase pathway was then examined using pharmacological inhibition. Treatment with the SphK2-selective inhibitor ABC294640 or the SphK1-selective inhibitor PF-543 alone had no significant effect on invasion. In contrast, the dual SphK1/2 inhibitor SKI II significantly reduced invasion to approximately 28% of control. Combined treatment of ABC294640 with PF-543 also had no significant effect, whilst combining ABC294640 with SKI II reduced invasion to approximately 14% of control. This was comparable to treatment with methyl-β-cyclodextrin (MβCD), a cholesterol-depleting agent, that reduced invasion to approximately 6% of untreated control, which has been found previously to have a significant effect on *E. coli* K1 uptake^4^ (Fig. 1D).

To further investigate the role of SphK1, cells were transfected with wild-type SphK1-GFP, a constitutively active gain-of-function mutant (G113A), or a kinase-dead mutant (G82D)^10,11^. Expression of wild-type SphK1 had no significant effect on invasion relative to the transfection control. However, expression of the G113A gain-of-function mutant significantly increased invasion to approximately 160% of control. There appeared to be a trend for increased invasion in the kinase-dead G82D mutant produced to 125%, although this was not found to be statistically significant (Fig. 1E). Overexpression of SphK2, using two independent GFP-tagged constructs (AcGFP-SphK2 and pcDNA SphK2 WT), increased the relative number of infected GFP-positive cells from approximately 3 to 5, suggesting that SphK2 overexpression also enhances bacterial invasion (Fig. 1F).

Finally, siRNA-mediated knockdown of sphingosine kinases paradoxically increased *E. coli* K1 invasion relative to the non-coding siRNA control. SphK1 knockdown increased invasion to approximately 210%, whilst SphK2 knockdown and combined SphK1+K2 knockdown increased invasion to approximately 320% and 330% respectively, both with statistical significance (Fig. 1G). This apparently contradictory result relative to the inhibitor data in Fig. 1D may reflect a non-catalytic scaffolding or adaptor role for the SphK proteins, whereby their physical presence, independent of enzymatic activity, restrains bacterial invasion. Taken together, these data confirm that *E. coli* K1 exploits EGFR signalling and the sphingosine kinase pathway during invasion of HBMECs in our system, and further suggest that SphK proteins play a dual role, both enzymatic and structural, in regulating bacterial entry.

### Flotillin 1 associates with *E. coli* K1 during intracellular infection of HBMECs

The finding that SphK inhibition and cholesterol depletion both impair *E. coli* K1 invasion suggests a role for sphingolipid-enriched membrane microdomains in facilitating bacterial entry. A mechanistic link between flotillin and both the SphK/S1P pathway^8^, as well as between flotillin and EGFR itself^7^ has previously been found. Therefore we wanted to investigate whether flotillin might also be involved in *E. coli* K1 uptake during HBMEC infection.

Following infection of HBMEC cells for either 30 min or 2 hours, differential labelling of extra- and intracellular bacteria combined with flotillin 1 immunostaining revealed a dynamic pattern of flotillin 1 recruitment to bacteria over time (Fig. 2A).

**Figure 2.**
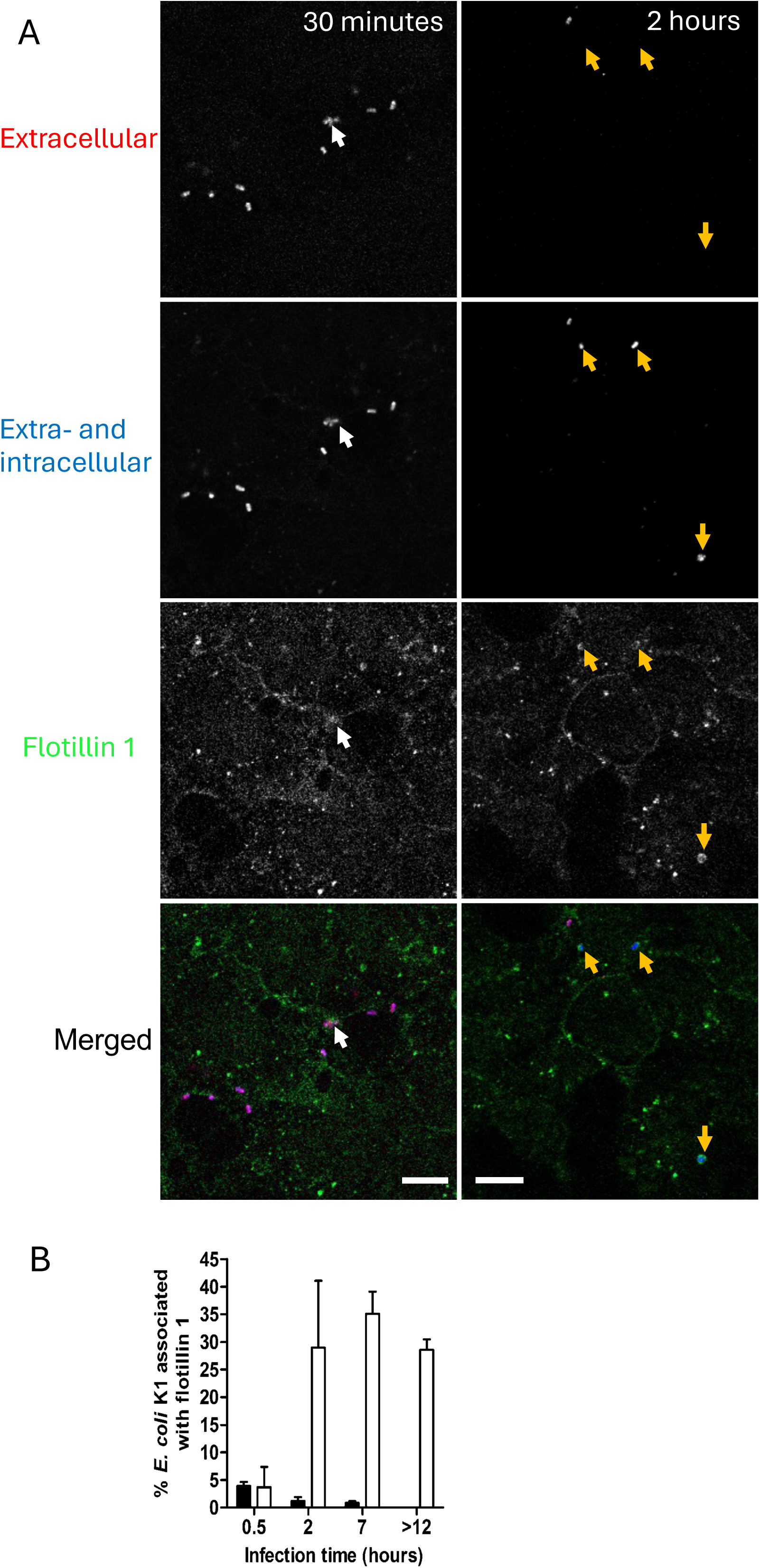
Association of flotillin 1 with extra- and intracellular E. coli K1 over time of infection. A. HBMEC were infected with E. coli K1 for the times indicated, fixed, and differential bacterial and flotillin 1 staining was performed to discern extra-from intracellular bacteria. Flotillin 1 is shown in green, extracellular bacteria in purple, intracellular bacteria in blue. A single confocal slice is shown. Flotillin 1 is associated with extracellular bacteria on the cell surface (white arrows), and concentrated around intracellular bacteria at the later time point (yellow arrows). Scale bars: 10µm. B. Quantification of bacteria-flotillin 1 association as in (A) during a timecourse of infection. Black bars, extracellular bacteria, white bars, intracellular bacteria. Data shown are the mean ± SEM from two independent experiments; ≥100 bacteria were counted per sample.

At 30 minutes post-infection, flotillin 1 was associated with bacteria at the cell surface (white arrows), consistent with an early interaction between the bacterium and flotillin-enriched plasma membrane microdomains. By 2 hours, the pattern had shifted markedly: flotillin 1 was now concentrated around intracellular bacteria (yellow arrows), with reduced surface-associated signal, indicating that flotillin 1 is recruited to bacteria as they are internalised and trafficked within the cell.

Quantification of bacteria-flotillin 1 association over a timecourse of infection confirmed this transition (Fig. 2B). At 30 minutes, a low number (approx. 5%) of extracellular bacteria associate with flotillin at the cell surface and this diminishes over time. Fewer than 5% of extracellular bacteria were associated with flotillin 1 at 30 min but this association increased substantially from 2 hours onwards, reaching approximately 30% at 2 hours, approximately 35% at 7 hours, and remaining elevated at approximately 29% beyond 12 hours. Together these data demonstrate that flotillin 1 is progressively recruited to *E. coli* K1 following internalisation, consistent with a role for flotillin-enriched endosomal compartments in intracellular bacterial trafficking, and providing a mechanistic link between the flotillin microdomain pathway and the SphK/S1P/EGFR axis identified above.

### Flotillin restricts *E. coli* K1 invasion in a cholesterol-dependent manner

The association of flotillin 1 with intracellular bacteria (Fig. 2) raised the question of whether flotillin promotes or limits bacterial entry. To test this, we manipulated flotillin function using genetic knockout, siRNA depletion and overexpression.

Caveolin-1 was included as a reference control, since we have previously shown that up- or down-regulation of caveolin does not significantly alter *E. coli* K1 invasion^4^.

Mouse lung endothelial cells (MLEC) lacking flotillin 1 showed markedly increased *E. coli* K1 invasion relative to wild-type cells, rising from approximately 0.003 to 0.05 cfu per cell (Fig. 3A). In contrast, caveolin-1 knockout MLEC did not differ from wild type, in agreement with our previous findings and indicating that the effect is specific to flotillin rather than a general consequence of disrupting raft-associated proteins. Consistent with this, overexpression of caveolin-1-GFP in HBMEC had no effect on the number of infected cells relative to the LDLR A18-GFP transfection control, whereas overexpression of flotillin 1-GFP reduced infection to approximately 22% of control and flotillin 2-GFP almost completely abolished it (Fig. 3B). Flotillin therefore appears to act as a negative regulator of bacterial invasion, with its loss enhancing uptake, and its overexpression suppressing it.

**Figure 3.**
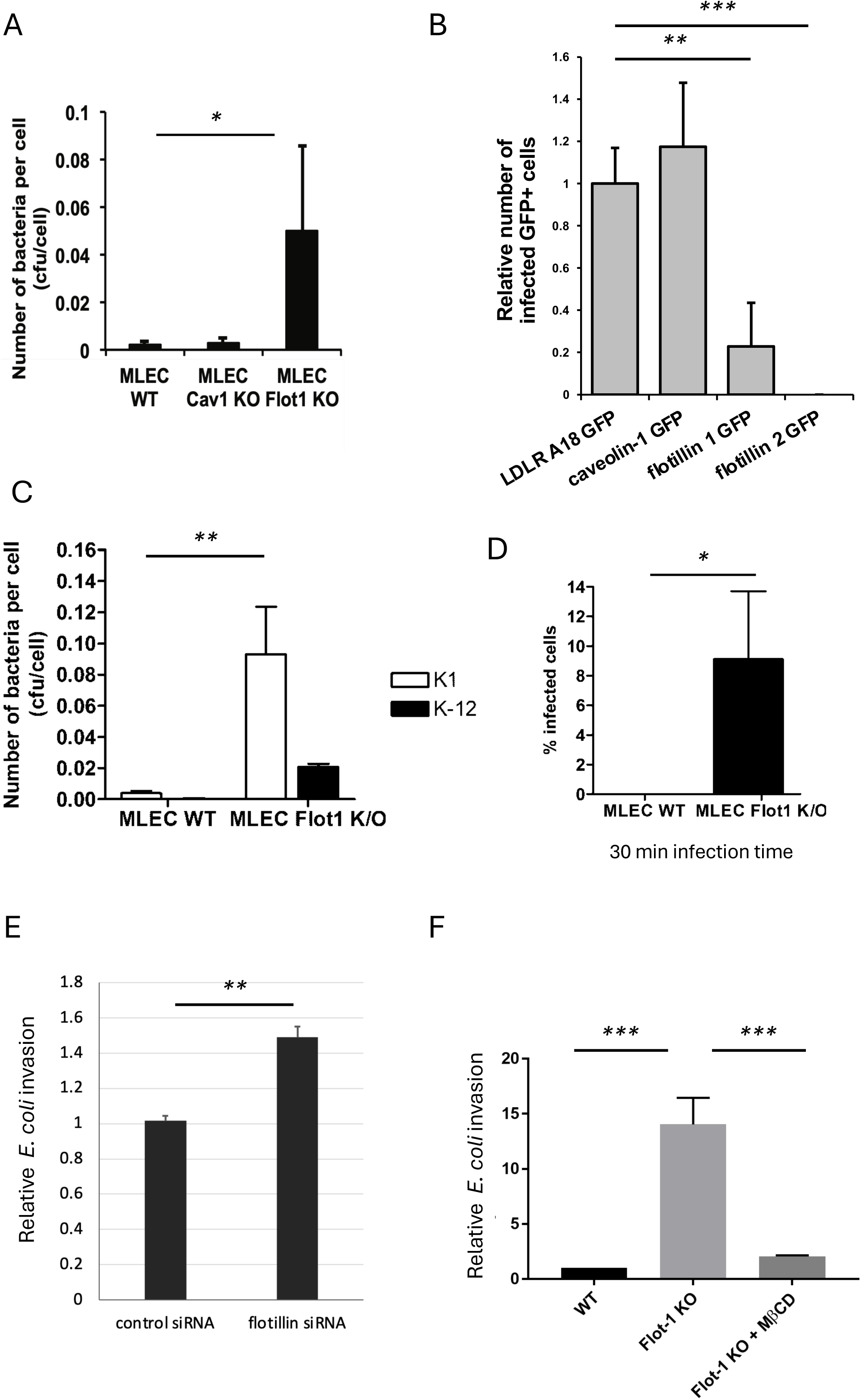
Manipulation of flotillin function alters *E. coli* K1 invasion. A. Mouse lung endothelial cells (MLEC) isolated from wild type, caveolin-1 knockout and flotillin-1 knockout mice were infected with *E. coli* K1 for 2 hours, then bacterial invasion analysed by cfu assay. B. HBMEC were transiently transfected with GFP chimaeras as shown. LDLR A18 GFP was a transfection control. GFP-expressing cells were scored for the presence of intracellular *E. coli* K1 following infection for 2 hours. C. MLEC from wild type or flotillin-1 knockout mice were infected with either *E. coli* K1 or non-pathogenic *E. coli* K-12 for 2 hours, then bacterial invasion analysed by cfu assay. D. MLEC isolated from wild type or flotillin-1 knockout mice were infected with *E. coli* K1 for 30 min, then fixed and differential bacterial staining was performed. Cells containing intracellular bacteria were counted, with ≥40 cells counted per sample. E. Uptake of *E. coli* K1 for 2 hours following siRNA depletion of flotillin-1 analysed by cfu assay. F. MLEC from wild type or flotillin-1 knockout mice were incubated with *E. coli* K1 ± methyl-β-cyclodextrin as shown. Data shown are mean ± SD from >3 independent experiments with triplicate samples. * *p*<0.05, ** *p*<0.01, *** *p*<0.005

We next asked whether this restriction is specific to the pathogenic *E. coli* K1 strain. Flotillin 1 knockout MLEC showed increased uptake of both *E. coli* K1 (∼0.09 cfu per cell, compared with ∼0.004 in wild type) and the non-pathogenic *E. coli* K-12 (∼0.02 cfu per cell, compared with near-undetectable levels in wild type), although the increase was considerably greater for *E. coli* K1 (Fig. 3C). This suggests that flotillin acts as a general barrier to bacterial uptake, with the pathogenic strain better able to exploit its absence. One possible reason for the increased bacterial count could be that they have increased replication once inside the flotillin knockout cells. To check whether the increas might be due to proliferation or whether due to invasion, uptake of bacteria at only a 30 min timepoint was investigated in the knockout cells. The effect was evident early in infection, with approximately 9% of flotillin 1 knockout cells contained bacteria after 30 min, whereas no infected wild-type cells were detected (Fig. 3D). Acute depletion of flotillin by siRNA in HBMEC similarly increased *E. coli* K1 invasion by approximately 50% relative to control siRNA (Fig. 3E), confirming that the phenotype is replicated across knockdown methods as well as between MLEC primary cells and HBMEC lines.

Finally, we tested whether the hyperinvasive phenotype depends on membrane cholesterol. Utilising cholesterol depletion with MβCD as in Fig. 1D, the increased uptake of *E. coli* K1 in flotillin 1 knockout cells, which showed an approximately 14-fold increase in invasion relative to wild type, was strongly suppressed by MβCD treatment, reducing to approximately 2-fold (Fig. 3F). Thus, the enhanced invasion in the absence of flotillin remains dependent on intact cholesterol-rich membrane domains.

### Loss of flotillin 1 increases penetration of the blood-brain barrier *in vivo*

To determine whether the restrictive role of flotillin observed in cultured endothelial cells (Fig. 3) is relevant *in vivo*, we used a neonatal mouse model of *E. coli* K1 infection. Wild-type and flotillin 1 knockout pups from asynchronous litters were infected through intraperitoneal injection at 5-7 days of age, and bacterial burdens in blood and brain were determined once animals became moribund.

Flotillin 1 knockout neonates had significantly lower bacterial numbers in the blood than wild-type littermates, with median counts of approximately 2 × 10⁶/ml compared with 3.5 × 10⁶/ml in wild type (Fig. 4A). Despite this reduction in bacteraemia, bacterial numbers in the brain were comparable between genotypes, at approximately 1-2 × 10⁵/ml in both groups (Fig. 4A). Because the blood is the source of bacteria that reach the brain, brain burden is best interpreted relative to bacteraemia. The brain-to-blood ratio was therefore significantly higher in flotillin 1 knockout mice than in wild-type mice, with a mean log₁₀ ratio of approximately 0.8 compared with 0.5, equivalent to an approximately two-fold increase in the efficiency with which circulating bacteria might be traversing to the brain (Fig. 4B). Thus, although flotillin 1 knockout animals carried fewer bacteria in the circulation, the same number reached the brain, indicating enhanced bacterial translocation across the blood-brain barrier. These findings are consistent with the increased invasion of flotillin-deficient endothelial cells *in vitro* and support a role for flotillin as a barrier to *E. coli* K1 entry into the central nervous system.

**Figure 4.**
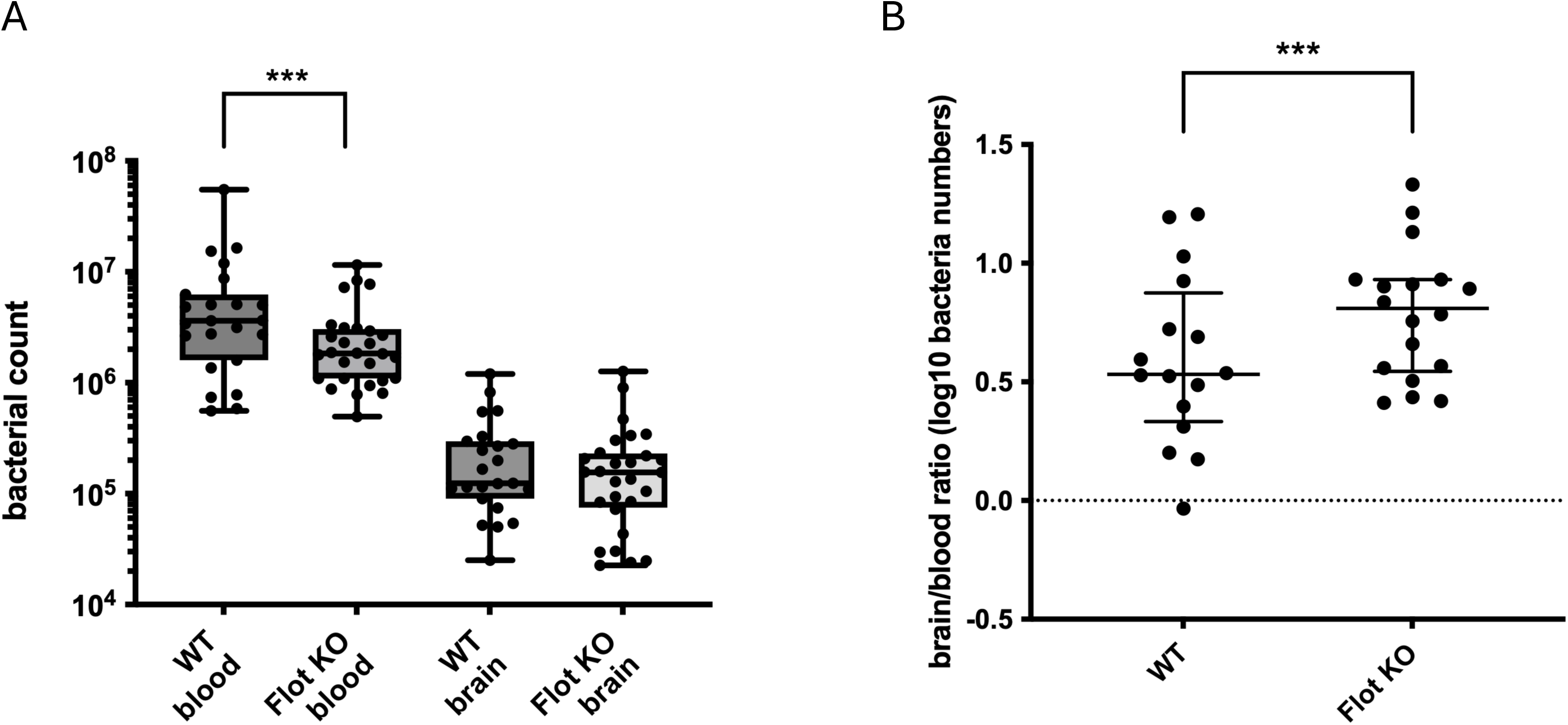
*In vivo* infection model shows increased invasion across the BBB in *flotillin 1^-/-^* mice. A. Bacterial count in tissues from moribund WT or *flotillin 1^-/-^* neonates. Asynchronous litters were injected i.p. with 5x10^4^ *E. coli* K1 at 5-7 days. B. Blood:brain ratio measurements from bacterial cfu were compared. Sample sizes ≥ 15 animals. *** *p*<0.005

## Discussion

Our data place flotillin 1 in a pivotal role at membrane-level control of *E. coli* K1 entry into brain endothelium. Our own and others’ data has demonstrated that invasion depends on EGFR, on sphingosine kinase activity and on cholesterol-rich membrane domains to enable macropinocytic uptake of the bacteria^3,4^. Flotillin 1 accumulates around intracellular bacteria suggesting a role in uptake and/or transcytosis. In contrast to caveolin, another protein that organises cholesterol-rich membrane domains, which has no effect on bacterial uptake, loss of flotillin markedly increases invasion *in vitro* and raises the brain-to-blood ratio of bacteria *in vivo*. We propose that *E. coli* K1 exploits a pro-invasive signalling cascade, while flotillin-dependent membrane organisation restrains the same process.

### A pro-invasion signalling cascade

Wang et al. showed that OmpA, FimH and NlpI activate SphK2 in HBMEC, and that the resulting S1P acts through S1PR2 to activate EGFR, both directly and by inducing HB-EGF expression and shedding^3^. Activated EGFR then recruits c-Src, driving actin rearrangement and uptake. Our data support the EGFR and SphK segments of this cascade in an independent system.

We found that overexpressing wild-type EGFR doubled invasion, and the Y1045F activated mutant, which evades downregulation^15^, tripled invasion. Erlotinib, an EGFR inhibitor, halved invasion. Since surface EGFR fell after an hour of infection in our study, but total EGFR does not change during infection^3^ this seems consistent with activation-driven internalisation. However since Wang et al demonstrated that EGFR inhibition does not alter bacterial adhesion^3^, and a host of other receptors are engaged in the process, it seems that EGFR must act indirectly.

For the sphingosine kinases, SKI II reduced invasion while the gain of function SphK1 G113A mutant increased it, and SphK2 overexpression similarly increased the number of infected cells. Together these suggest that a greater S1P-generating capacity might help to promote entry.

### Flotillin as a restraint on entry

Loss of flotillin 1 increased invasion in knockout MLEC and after siRNA depletion in HBMEC, whereas overexpressing flotillin 1 or 2 significantly abrogated uptake. Caveolin served as a reference control: neither its loss nor its overexpression altered invasion, as shown in our previous work^4^. The restriction by flotillin that is alleviated in the knockout is not wholly specific to *E. coli* K1, since *E. coli* K-12 uptake also rose in knockout cells, although the effect was larger for *E. coli* K1. Imaging at 30 minutes showed infected knockout cells but none in wild type, indicating flotillin’s effect on entry rather than only on survival, despite the clear colocalization with bacteria once inside the cell. *In vivo*, *flotillin 1^-/-^* neonates had lower bacteraemia but similar brain burdens, giving a higher brain:blood ratio.

Because animals were sampled when moribund, this suggests that knockouts reach a comparable CNS burden from a lower circulating load, i.e. more efficient translocation.

The decreased blood bacteraemia is particularly interesting as an effect and how this might arise in flotillin-deficient mice seems strange. These knockout mice have impaired neutrophil recruitment^16^ and, since neutrophils are normally a primary defense to destroy bacteria, it therefore might be expected there would be increased bacteraemia as an effect as bacteria should be able to multiply unchecked.

Alternatively since endothelial uptake of bacteria is increase, maybe the bacteria may be taken up by endothelia throughout the bloodstream, could contribute to altered bacteraemia. Additionally, because flotillin microdomains reside on late endosomes and lysosomes^8^, the intracellular association between the bacteria and flotillin may reflect maturation of bacteria-containing vacuoles. The cfu assays measure net intracellular bacteria, so flotillin may restrict both entry and survival.

### How might flotillin restrain invasion?

Loss of flotillin did not remove the cholesterol requirement. MβCD reduced the hyperinvasion of knockout cells by approximately 85%, so the route used in the absence of flotillin remains cholesterol-dependent, consistent with the macropinocytic pathway we described previously^4^.

Flotillins form their own cholesterol-dependent microdomains that support a clathrin-independent endocytic pathway, promote membrane curvature and engage the actin cortex^5,16,17^.

The increase in bacterial uptake upon loss of flotillin might be explained by a number of routes, such as if flotillin microdomains sequester cholesterol away from sites of ruffling, then loss of flotillin increases the membrane available for micropinocytosis. Alternatively if flotillin loss weakens actin organisation, it might make the plasma membrane more permissive to remodelling by the bacteria.

A further possibility is that flotillin regulates EGFR itself, since flotillins associate with EGFR and influence its clustering and signalling^7,18^, and flotillin-1 depletion increases EGFR expression and signalling^19^. EGF can also induce tyrosine phosphorylation on flotillin-2^20^ and Fyn regulates flotillin endocytosis^21^. Infection induced EGFR and Src activity might therefore phosphorylate flotillin and facilitate internalisation with bacteria.

EGFR uptake can use clathrin-mediated endocytic pathways or micropinocytosis^6^. EGF is a potent activator or macropinocytosis and to avoid unchecked signaling EGFR must be regulated through internalisation. Without flotillin, EGFR remains stimulated but not internalised and this leads to enhanced macropinocytosis^7^. This would fit, then, in loss of flotillin, that EGFR activity would be elevated, enabling hyperinvasion by the bacteria.

### How does flotillin regulate S1P signalling

Riento et al. showed that flotillins bind sphingosine, recruit it to membranes and maintain S1P levels, with S1P reduced in knockout tissues (brain included) but not in serum^8^. Combined with Wang et al.^3^, this predicts that flotillin loss should reduce S1P-driven invasion, yet we observe the opposite. It may be that S1P activates EGFR and promotes entry independently of flotillin, while alternatively S1P may be important for membrane barrier integrity. However since we found that both flotillin depletion and SphK siRNA both increase invasion it seems they must be in the same pathway.

### Why does SphK knockdown increase invasion?

SphK1 and SphK2 siRNA increased invasion (significantly for SphK2 and the combination), whereas inhibitors decreased it, wild-type SphK1 overexpression had no effect and only the gain-of-function mutant increased entry. This seems counterintuitive. But since inhibitors such as SKI II block only the active site whilst leaving the protein intact, in contrast to sirRNA-mediated depletion which removes the protein entirely, potentially abolishing protein interactions or structural functions that might normally limit bacterial entry.

Consistent with this, the kinase-dead G82D mutant also showed a trend towards increased invasion despite the loss of catalytic activity, suggestion that the presence of the protein may itself influence invasion, and SphK1 has been implicated in forming endocytic structures^22^. Further, the inhibitors have some cross effects with SKI II also possibly impacting ceramide pathway^23^, and ABC29460 might even cause increased S1P levels^24^. Therefore the SphK proteins might play a dual role, both enzymatic and structural, in regulating bacterial entry.

### Relationship to the *sphK2^-/-^* phenotype

Wang et al. found reduced brain penetration in *sphK2^-/-^*mice^3^, which appears to conflict with our data. This may be accounted through differences in the infection model where their brain counts were 1 hour after intracardiac injection, and in this investigation they were taken at a moribund endpoint, much later. Furthermore *sphK2^-/-^* mice have elevated systemic S1P^25^, which could strengthen S1PR1-mediated barrier function independently of endothelial SphK2 activity. The flotillin knockout instead lowers S1P in tissues, including brain^8^, therefore the two models therefore might cause opposing S1P effects in different compartments.

Overall our data suggest that *E. coli* K1 hijacks an S1P-EGFR-Src cascade that flotillin-dependent membrane organisation normally restrains. Targeting this axis could provide strategies to limit brain penetration of these pathogenic bacteria. This would create complementary approaches to supplement antibiotic treatments, which may reduce the long-term neurological sequelae in neonate patients.

## Notes

### Competing Interest Statement

The authors have declared no competing interest.

### Funding

- Bloomsbury Colleges, London UK

